# Phosphoproteomic profiling reveals post-translational dysregulation in Huntington’s disease patient-derived neurons

**DOI:** 10.1101/2025.10.24.684372

**Authors:** Lea Danics, Chandramouli Muralidharan, Ágnes Varga, Melinda Rezeli, Jeovanis Gil, Anna A. Abbas, Ádám Pap, Andrew S. Park, Marcell Cserhalmi, Ármin Sőth, Dorina Jamniczky, Roland Zsoldos, Roger A. Barker, Gergely Róna, Janelle Drouin-Ouellet, György Markó Varga, Zsuzsanna Darula, Karolina Pircs

**Author notes:** Correspondence to: Karolina Pircs Semmelweis University, Institute of Translational Medicine 1094, Budapest Hungary. These authors have contributed equally to this work (Lea Danics, Chandramouli Muralidharan).

## Abstract

Huntington’s disease (HD) is a fatal neurodegenerative disorder caused by a CAG repeat expansion in the *Huntingtin* (*HTT*) gene. Although transcriptomic and proteomic changes have been characterized in patient-derived neurons, the contribution of post-translational modifications (PTMs), such as phosphorylation, remains poorly understood. Here, we present the first phosphoproteomic analysis by mass spectrometry (P-MS) of human induced neurons (iNs) directly reprogrammed from HD patient fibroblasts. We identified 177 phosphopeptides with significantly altered abundance in HD-iNs, mapping to phosphoproteins associated with key signaling pathways known to be affected in HD, such as splicing and autophagy. By integrating P-MS data with previously published proteomic and transcriptomic data from the same donors, we identified distinct subsets of ON-OFF phosphopeptides that exhibited a complete loss of phosphorylation in either HD- or Ctrl-iNs, without corresponding changes at the RNA or protein level. An exception was MXRA8, previously described in glial cells as a mediator of blood-brain barrier integrity and astrocyte-mediated neuroinflammation. This protein showed increased protein abundance despite the absence of phosphorylation in HD- iNs, suggesting a compensatory mechanism - a pattern also observed in human post mortem cortical HD tissue. Additionally, MXRA8 showed altered protein-protein interactions with lysosomal and metabolic regulators in HD-iNs, highlighting its potential role in autophagy impairment as well as in neurovascular dysfunction. These findings uncover a distinct layer of post-translational dysregulation in HD, suggesting that phospho-switch proteins such as MXRA8 may be candidate effectors of pathology and thus site-specific phosphorylation loss may contribute to impaired signaling and proteostasis in human HD neurons.

## INTRODUCTION

Huntington’s disease (HD) is an autosomal dominant genetic disorder characterized by age-dependent progressive neurodegeneration predominantly in the striatum and in the cortex of the brain [1, 2]. Although HD is considered a rare disorder, studies show consistent increase in HD incidence and prevalence over the past decades, posing a growing burden on families, healthcare systems and society [3–6]. HD manifests with progressing cognitive impairment, involuntary movements, psychiatric, metabolic and sleeping problems and eventually death of the patient typically within about 20 years after the symptoms first appear [1, 7]. Despite decades of research, no curative treatment is currently available.

The disease is caused by an expanded CAG repeats in one of the alleles of the *Huntingtin (HTT)* gene, translated to a mutant HTT (mHTT) protein containing an abnormally long poly-glutamine (polyQ) tract near the N-terminus [2]. This mHTT protein exhibits toxic gain-of-function properties and has a greater tendency to misfold, fragment, and form aggregates when compared to the wild-type protein [1, 2]. However, beyond the toxicity of mHTT in HD, numerous pathways, associated with autophagy, intracellular trafficking, energy metabolism, synaptic function and protein homeostasis, are disrupted, leading to the pathophysiology and progression of the disease [1, 8]. A better understanding of the molecular changes involved in such pathways in HD could lead to alternate therapeutic interventions that target the affected pathways.

Protein phosphorylation, the most prevalent post-translational modification (PTM), is a critical player in cell signaling pathways and is known to be dysregulated in various neurodegenerative disorders. Additionally, dysregulation of phosphorylation has previously been shown to play a pivotal role in the pathology of HD [9–16]. However, there is a lack of in-depth characterization of phosphorylation events in HD, which could offer deeper insights into the altered signaling pathways and help discover potential new therapeutic targets.

Studying HD remains challenging, as it is a human-specific disorder and is strongly influenced by aging, features that are difficult to replicate in conventional cellular or animal models [17–20]. In this study, we performed high-throughput phosphoproteomic analysis by mass spectrometry using a previously established induced neuronal (iN) model derived from HD patient fibroblasts with age- and sex-matched controls, that retains the age-related epigenetic characteristics of the donor [21, 22]. When integrated with proteomic data from the same cohort in our previous study [18], we observed striking alterations in the phosphoproteome of HD-iNs compared to their proteome profiles, suggesting that PTMs may play a significant role in HD pathology. We identified 46 phosphosites across 43 proteins that were phosphorylated exclusively in either control iNs or HD-iNs, referred to as ON-OFF and OFF-ON phosphoproteins, indicating disrupted phosphorylation regulation of key proteins and pathways.

Strikingly, only one ON-OFF protein in HD-iNs, the matrix remodeling associated protein 8 (MXRA8), showed a complete lack of phosphorylation at a specific site, coupled with a significant increase in abundance at the protein level, indicating a possible compensatory effect to MXRA8 dysfunction. Using a custom-made phospho-specific antibody, we validated this P-MS and MS phenotype of MXRA8 in HD-iNs. Additionally, we identified several candidate kinases involved in regulating MXRA8 phosphorylation and confirmed dysregulation of some of these by western blotting (WB). Consistent with these findings, immunohistochemical analysis of post mortem HD cortical tissue revealed neuronal MXRA8 expression with an apparent increase in MXRA8 signal intensity compared to controls, supporting a similar trend in the human HD brain. Lastly, we found altered MXRA8 binding partners through co-immunoprecipitation (co-IP) coupled with MS, which revealed a novel role in neuronal autophagy regulation and neurovascular function [23].

This study therefore presents the first open-access phosphoproteomic landscape of human HD-iNs. By combining our previously published proteomic analysis with this new phosphoproteomic data using the same HD samples, we have provided a multiomic approach to better understand HD pathophysiology [22]. Detection of ON-OFF proteins such as MXRA8 highlights new mechanistic processes in HD pathology, potentially leading to the identification of novel therapeutic targets and underscoring the translational potential of PTMs.

## METHODS

### Fibroblast lines

In this study we used dermal fibroblast samples from a previously published cohort of seven HD patients and seven age-matched, non-genetically related healthy individuals, as controls (Ctrl) (Table 1) [22]. Fibroblasts were obtained via skin biopsy, as described previously [21]. The skin biopsy samples were obtained from the Huntington’s Disease Clinic at the John van Geest Centre for Brain Repair (Cambridge, UK) and the Fondazione IRCCS, Instituto Neurologico Carlo Besta (Milan, Italy) and used under local ethical approvals (REC 09/H0311/88, IV-2625-1/2021/EKU, IV-1029-1/2022/EKU). The CAG repeat length was defined by Sanger sequencing for both alleles (Laragen Sanger Sequencing Services). The age of cohort ranged between 27 and 66 years old. CAG repeat length of the *mHTT* allele in the HD patients was between 39 and 45 repeats [24]. The healthy controls had CAG repeats less than 24 (Table 1).

**Table 1.** Demographic summary of Ctrl and HD patients.

| ID | Age | Sex | CAG repeats | Age at onset of HD |
| --- | --- | --- | --- | --- |
| HD1 | 28 | M | 15/39 | premanifest |
| HD2 | 31 | M | 20/45 | 33 |
| HD3 | 43 | M | 17/42 | 38 |
| HD4 | 43 | M | 19/44 | 36 |
| HD5 | 47 | M | NA/40 | premanifest |
| HD6 | 53 | M | 19/42 | premanifest |
| HD7 | 59 | M | 16/39 | 33 |
| C1 | 27 | M | 17/17 | - |
| C2 | 30 | M | 19/24 | - |
| C3 | 52 | F | 19/23 | - |
| C4 | 54 | F | 15/20 | - |
| C5 | 61 | M | 17/23 | - |
| C6 | 61 | F | 17/17 | - |
| C7 | 66 | M | 24/24 | - |

### Cell culture

Fibroblasts were kept in Glutamax supplemented high-glucose Dulbecco’s Modified Eagles Medium (DMEM) (Gibco) containing 10% foetal bovine serum (FBS) (Euroclone or Hyclone) and 1% Penicillin-Streptomycin (Gibco). Cells were passaged at 80 - 90% confluency following the protocol described previously [22].

### Lentiviral production and neuronal conversion

Third-generation lentiviral vectors were produced following a previously described protocol [25]. Virus titration was performed by RT-qPCR and the titer determined as previously described [25]. Virus titers ranged between 2.70 × 10^8^ and 1.86 × 10^9^.

For the direct conversion of dermal fibroblasts into induced neurons the previously published LV.U6.shREST1.U6.shREST2.hPGK.BRN2.hPGK.ASCL1.WPRE all-in-one transfer vector was used [21]. The construct available from the plasmid repository, contains two short hairpin RNAs (shRNA) targeting Repressing Element Silencing Transcription Factor (REST) 1 and 2 after a U6 promoter and transcription factors ASCL1 and BRN2 after the non-regulated ubiquitous phosphoglycerate kinase (PGK) promoter and a Woodchuck Hepatitis Virus (WHP) Post-transcriptional Regulatory Element (WPRE).

Fibroblasts were plated at a density of ∼26 000 cells/cm^2^ on Nunc Delta surface treated 24-well plates, (Thermo Scientific) pre-coated with poly-L-ornithine (Sigma), Fibronectin (Thermo Fischer) and Laminin 111 (Biolamina) or tissue-culture treated T25 and T75 flasks (Thermo Scientific) pre-coated with 0.1% gelatine (Sigma) [22, 26]. Twenty-four hours later the fibroblasts were transduced with the all-in-one lentiviral vector at MOI 10 or 20 and incubated for 72 hours. Neural conversion was continued with growth factors and small molecules containing NDiff medium (Takara Bio) as previously described [21, 22]. iNs were harvested or fixed on day 28 for further experiments.

### Phosphoproteomic analysis by mass spectrometry

iNs converted in T75 flasks (600,000 fibroblasts plated for conversion per sample) were dissociated as previously described and prepared for phosphoproteomic analysis as follows [22]. The cells were carefully washed off and collected in a tube with Accutase and spun at 400 x g for 5 minutes. The supernatant was discarded, and the pellets were washed three times with DPBS. After the final wash, the supernatant was aspirated, and the pellets were frozen on dry ice and stored at -80 °C until use.

The cell pellets were resuspended in 200 µL lysis buffer (50 mM DTT, 2 %SDS, 100 mM Tris pH = 8.6), rested for 1 min on ice and sonicated (20 cycles: 15 seconds on/off; Bioruptor plus model UCD-300, Diagenode). Reduction and alkylation of disulfide bridges was performed by incubating the samples at 95 °C for 5 minutes, followed by the addition of iodoacetamide to a final concentration of 100 mM with and incubation for 20 minutes at room temperature in the dark.

Samples were processed using S-Trap Mini Spin Columns (ProtiFi, USA) according to the manufacturer’s instructions. Briefly, samples were acidified by adding phosphoric acid to a final concentration of 1.2%, 7 volumes of binding buffer (90% MeOH, 100 mM TEAB, pH = 7.1) was added to the samples, which were then transferred to the S-Traps, and spun at 4000 x g for 30 seconds. The trapped proteins were washed three times with the binding buffer. Protein digestion was performed by adding trypsin (Promega Biotech AB) 1:50 (enzyme:protein ratio) in 125 µL of 50 mM TEAB and incubating for 16 hours at 37 °C. Peptides were eluted with 0.2% of aqueous formic acid and 0.2% of formic acid in 50:50 water:acetonitrile. Following speed vacuum concentration, peptides were dissolved in 0.1% TFA and quantified with the Pierce Quantitative colorimetric peptide assay (Thermo Fisher Scientific). 50 μg of peptides were subjected to metal ion affinity chromatography (IMAC) enrichment using Fe(III)-NTA cartridges on the Agilent AssayMAP Bravo platform (Agilent Technologies), as previously described [27]. The enriched phosphopeptides were resuspended in 20 μL of 2% ACN with 0.1% TFA after speed vacuum concentration, and 15 μL was injected for each sample on the nanoLC-MS/MS system.

NanoLC-MS/MS analysis was performed on a Dionex Ultimate 3000 RSLCnano UPLC system coupled to a Q-Exactive HF-X mass spectrometer (Thermo Fischer Scientific).

Phosphopeptides were first trapped on an Acclaim PepMap100 C18 trap column (3 µm, 100 Å, 75 µm i.d. × 2 cm, nanoViper) and separated on an EASY-spray RSLC C18 analytical column (2 µm, 100 Å, 75 µm i.d. × 25 cm) using a non-linear gradient. Solvent A consisted of 0.1% formic acid (FA), while solvent B contained 80% ACN with 0.08% FA. The flow rate was set to 0.3 μl/min, and the column temperature was 45 °C. The gradient was initiated at 4% solvent B, ramped to 27% over 120 minutes, increased to 45% over the next 15 minutes, then rapidly increased to 98% within 1 minute, and held at 98% for an additional 5 minutes. A top 15 DDA method was applied, where MS1 full scans were acquired with a resolution of 120,000 (@ 200 m/z), a target AGC value of 3E+06, and a maximum injection time of 50 milliseconds, using a mass range of 375-1500 m/z. The 15 most intense peaks were fragmented with an NCE of 25. MS2 scans were acquired with a resolution of 60,000, a target AGC value of 1E+05, and maximum IT of 120 milliseconds. The ion selection threshold was set to 7.2E+03 and the dynamic exclusion to 30 seconds.

Protein identification and label-free quantification was performed using Proteome Discoverer v2.2 (Thermo Fisher Scientific) with SEQUEST HT as the search engine and a human protein database downloaded from UniProt on 2019-01-15. Trypsin was specified as the protease, allowing for up to two missed cleavages. The mass tolerance was set to 10 ppm for MS1 and 0.02 Da for MS2. Carbamidomethylation of cysteine was set as a static modification, while methionine oxidation, phosphorylation on serine, threonine and tyrosine, and protein N-terminal acetylation were included as dynamic modifications. Phosphorylation site scoring was performed using the ptmRS algorithm, applying a site probability threshold of >75. Peptides and corresponding proteins were identified at a 1% false discovery rate (FDR).

All phosphopeptide intensities were log2-transformed and median-centered per sample to correct for differences in sample loading. Technical replicates were averaged, and only peptides quantified in at least four of seven samples in at least one group were retained, yielding 4,363 phosphopeptides corresponding to 1,493 different proteins. Additionally, phosphopeptide intensities were normalized to the corresponding protein abundances using matched global proteome data [22]. Specifically, each phosphopeptide was mapped to its parent protein in the global dataset, and normalization was performed by subtracting the log2-transformed protein intensity from the log2-transformed phosphopeptide intensity (log2[phosphopeptide] − log2[protein]). The resulting values represent phosphorylation changes independent of protein expression. Prior to this correction, missing protein intensities in the global proteome were imputed from the lower tail of the intensity distribution to avoid artificially inflating the phospho-to-protein differences. Principal component analyses were performed in Perseus v1.6.5.0.

### Phosphoproteomic Gene-centric analysis

Using a combination of Shapiro-Wilk and an unpaired t-test on R v4.3.1 [28] differential abundance analysis was performed on the phosphopeptides, comparing HD-iNs with Ctrl-iNs. All phosphopeptides with a Shapiro-Wilk p > 0.05, or with a mean abundance of 0 in either Ctrl-iNs or HD-iNs were considered for unpaired t-test. Subsequently, those with a t-test p < 0.05 were considered significant. Further, pathway enrichment analysis was performed on the phosphoproteins corresponding to the significantly differentially abundant phosphopeptides, using STRING-db v12.0[29] on the corresponding website, with a background combining all the proteins detected in the phosphoproteomic data and those in the proteomic data from Pircs et al [22].

### Phosphosite Enrichment Analysis

Phosphosite annotated peptide sequences across the differentially abundant phosphopeptides, along with their log2-fold change were uploaded on to the PSEA webtool of Kinase Library on the PhosphoSitePlus website [30–32]. The results were downloaded as tab-delimited files and further analysed using R v4.3.1. All kinases with a p-value < 0.05 and those previously identified in the proteomic data from Pircs et al. [22], were considered significant and inspected further.

### Kinase Prediction

The phosphosites associated with the ON-OFF phosphopeptides were each manually searched for the PhosphoSitePlus website, and using the Kinase Library’s kinase prediction webtool, kinases were predicted [30–32]. The kinase prediction results for each phosphosite were downloaded as tab-delimited files and further analysed using Rv4.3.1. All kinases with a site percentile score greater than 90 and previously identified in the proteomic data from Pircs et al. [22], were considered significant and inspected further.

### Immunocytochemistry

Immunocytochemistry (ICC) on iNs was performed using a previously described protocol [21, 22]. Briefly, on day 28 iNs were fixed with 4% PFA (pH = 7.4) at room temperature for 10 minutes. After washing with DPBS, iNs were permeabilized with 0.1% TritonX in DPBS for 10 minutes and then blocked for 30 minutes with a blocking solution of 5% donkey serum in DPBS. After blocking, primary antibodies diluted in blocking solution were added to the cells and incubated overnight at 4°C (Table 2). After removing the primary antibodies, the cells were washed twice with DPBS and then incubated for 2 hours at room temperature in the dark with fluorophore-conjugated secondary antibodies diluted in blocking solution (Table 2). Secondary antibodies were removed by washing the cells with DPBS, then DAPI staining was applied and incubated for 15 minutes at room temperature, in the dark. DAPI staining was removed, and cells were washed with DPBS before high-content automated microscopy analysis.

**Table 2.** List of antibodies used for ICC and IHC.

| Antibody | Conc. | Species | Company | Cat# | RRID |
| --- | --- | --- | --- | --- | --- |
| TAU (clone HT7) | 1:500 | Mouse | Thermo Scientific | MN1000 | AB_2314654 |
| MXRA8 | 1:100 | Rabbit | Abcam | Ab185444 | N/A |
| MAP2 | 1:1000 | Chicken | Abcam | AB5392 | AB_2138153 |
| MAP2 | 1:1000 | Guinea Pig | Synaptic Systems | 188004 | AB_2138181 |
| Alexa Fluor® 488 AffiniPure Donkey IgG | 1:200 | Mouse | Jackson | 715-545-150 | AB_2340846 |
| Alexa Fluor® 647 AffiniPure Donkey IgG | 1:200 | Rabbit | Jackson | 711-605-152 | AB_2492288 |

### High-content automated microscopy

A CellInsight CX5 High-content Automated Screening platform was used for microscopic analysis of the immunostained iNs. By this method a fast and unbiased analysis can be performed. For the identification of DAPI^+^ and TAU^+^ cells, we used the target activation (TA) protocol of the CX5 HCS software. Images were acquired from 25 or 100 fields on 96-well plates using a 10× or 20× objective, respectively, and wells were included for further analysis, in which there were at least 50 valid neurons. The TA protocol defined DAPI^+^ cells by intensity, shape and area values. Border objects and DAPI^+^ cells which were aggregated and where single nuclei could not be segmented by the software, they were excluded. Within the DAPI^+^ cells, TAU^+^ cells were then defined based on average cell body fluorescent intensity and area. Purity of the iN culture was determined as the fraction of TAU^+^ cells of the total DAPI^+^ cells and conversion efficiency as the number of TAU^+^ cells over the starting number of fibroblasts used for that conversion [22].

For analyzing MXRA8 staining the neural profiling protocol was applied with spot detection using a 20× objective on 144 or 529 fields per each well of a 96-well plate or 24-well plate respectively, and wells were included for further analysis, in which there were at least 50 valid neurons. First, DAPI^+^ nuclei and TAU^+^ cells were identified as described above, and TAU^+^ cell bodies and neurites were defined as the region of interest (ROI). Then, the average number and area of different dots per cell in TAU^+^ cell bodies and neurites were determined. In addition, a separate spot detection analysis was performed in which the average number and size of MXRA8 dots were quantified specifically within the nuclei of TAU^+^ cells. Border objects and wells with less than 50% valid fields were excluded from analysis.

The fluorescent intensity, shape and area threshold settings were defined on randomly chosen fields of view from both groups of iNs. The accuracy of the settings was verified by a short pre-analysis on 10 images from each well which was validated by an independent researcher.

### Immunofluorescence on paraffin-embedded human brain sections

The Cambridge Brain Bank provided anonymous, 10-μm thick paraffin-embedded tissue sections from HD patients (N = 3) and age-matched controls (N = 2) known not to have any neurological or psychiatric disorders (Table 3). Cortical tissue was available for all cases. Demographic data was obtained from the Brain Bank. The pathological severity of HD was scored according to the Vonsattel grading system [33]. Brains sections were used under full local ethical approval (CERC-2025-7229; University of Montreal).

**Table 3.** Human post mortem brain tissue samples. (NA: not applicable, M: male, F: female)

| Brainbank ID | Age at death | Sex | Pathological grade |
| --- | --- | --- | --- |
| H682 | 40 | M | 4 |
| H720 | 68 | M | 4 |
| H725 | 58 | M | 4 |
| PT149 | 56 | M | NA |
| PT155 | 39 | F | NA |

Deparaffinized and rehydrated tissue sections were incubated overnight at 4°C with the following primary antibodies: Rabbit anti-MXRA8, guinea pig anti-MAP2 and chicken anti-MAP2. On the second day, after washing three times with PBS, Cyanine-conjugated secondary antibodies (1:200; Jackson ImmunoResearch Laboratories) were added and counterstained with DAPI (1:1,000; Sigma-Aldrich). To reduce tissue autofluorescence, sections were treated with an autofluorescence eliminator reagent (Sigma). Controls included staining after omitting the primary antibodies to distinguish tissue autofluorescence from specific antibody staining. Fluorescent images of 2048px by 2048px were acquired with a Nikon AX/AX R with NSPARC confocal microscope equipped with a 20x lens (PLAN APO 20x DIC M/N2).

### Custom phospho-antibody design

For phospho-MXRA8 protein identification we ordered a custom rabbit polyclonal phospho-peptide antibody from Biomatik (Ontario, Canada). The antibody recognizes the protein sequence including the phospho-S377 (pS377) phospho-site of MXRA8 protein. The sequence of the designed phosphopeptide is Cys-GGYEYSDQK(pS)GKSKGKD representing [367–384] residue of human MXRA8 containing an additional Cys at the N-terminus of the peptide for chemical conjugation purposes (pS represents the phosphorylated Ser). A non-phosphopeptide with the same sequence, with the exception of a non-phospho-S377 instead of pS377, was used as negative control. For antibody production a standard 70-day immunization protocol was used. From 2 immunized rabbits 1 ml immune serum was collected in total and the isolated antibodies were purified by affinity chromatography. The quality of the antibody was verified by ELISA measurement (serial dilution >1:32, 000; OD = 1).

### Western blot

For western blot (WB) experiments, 200,000 fibroblasts were converted into iNs in a 0.1% gelatine-coated T25 flask for each sample. iNs were harvested on day 28 by dissociating with Accutase (Corning) and then collected with HBSS (Gibco) and spun at 400 g for 5 minutes. The cell pellets were lysed with RIPA buffer (Sigma) containing a 4% complete protease inhibitor cocktail (PIC) (Roche) and 4% PhosSTOP phosphatase inhibitor (Roche). Cell lysates were collected and incubated on ice for 30 minutes followed by centrifugation at 10,000 × g for 10 minutes at 4 °C to sediment cellular debris. The supernatant was collected and stored at -80 °C until further use. Gel electrophoresis and semi-dry blotting was performed as described previously [34]. A STK10 WB was made following the manufacturer’s instructions: primary Ab was incubated for 1.5 h at RT, after washing, secondary Ab was incubated for 1 h at 37 °C. Primary and secondary antibodies were diluted in 5% milk blocking solution (Table 4). For blot detection the Immobilon Western Chemiluminescent HRP Substrate (Millipore) was used, blots were incubated for 5 minutes at room temperature in the dark followed by immediate visualization by Imager2Imager CHEMI Premium Gel Documentation System (VWR). Band intensity was quantified using Fiji software and corrected to the amount of beta-actin reference protein per each lane.

**Table 4.** List of antibodies used for WB.

| **Antibody** | **Conc.** | **Species** | **Company** | **Cat#** | **RRID** |
| --- | --- | --- | --- | --- | --- |
| MXRA8 | 1:1,000 | Rabbit | Abcam | Ab185444 | N/A |
| MXRA8 | 1:1,000 | Rabbit | Biomatik | N/A (custom made) | N/A |
| MXRA8-pS377 | 1:1,000 | Rabbit | Biomatik | N/A<br>(custom made) | N/A |
| MAPK9 | 1:1,000 | Rabbit | Cell<br>Signaling<br>Technology | 9258T | N/A |
| STK10 | 1:500 | Rabbit | Proteintech | 25471-1-AP | AB_2880096 |
| EEF2K | 1:1,000 | Rabbit | Invitrogen | PA5-22175 | AB_11153234 |
| Hrp Conjugated<br>antibody | 1:5,000 | Rabbit | Cytiva | NA9340 | AB_772191 |
| β-actin | 1:50,000 | Mouse | Sigma-Aldrich | A3854 | AB_262011 |

### Co-Immunoprecipitation (co-IP), mass spectrometry and data analysis

Immunoprecipitation experiments were performed as described previously [35]. Briefly, fibroblasts or iNs were collected into pre-chilled 1.5 ml tubes using Accutase for detachment. The cells were spun down at 300 × g for 5 min at 4 °C, then the pellet was washed with PBS and spun down again. The supernatant was removed, and the pellets were snap frozen and stored at -80 °C until further processing.

Next, the cells were lysed in lysis buffer (25 mM Tris.HCl pH 8.0, 150 mM NaCl, 0.2% NP-40, 10% glycerol, 1 mM EDTA, 1 mM EGTA, 2 mM MgCl2, 0.5 mM 1,4-Dithiothreitol (DTT) and 5 mM N-Ethylmaleimide (Sigma-Aldrich)) and containing 5 U/mL TurboNuclease (Accelagen)) for 30 min on ice aided with a 1 min burst of sonication (Joanlab UC30D). The insoluble fraction was removed by centrifugation (20,000 × g for 10 min at 4 °C). The collected supernatant corresponds to the soluble fraction. All the above buffers were supplemented with protease (Complete™ ULTRA, Roche) and phosphatase inhibitors (PhosSTOP™, Roche). Endogenous immunoprecipitations of MXRA8 were carried out for 2 h at 4°C using anti-MXRA8 antibody (Abcam, Table 4) mixed with Protein A/G Magnetic Beads (Pierce) at 4°C for 2 hours. Beads were extensively washed in lysis buffer and an elution was carried out with elution buffer (70 mM Tris HCl, pH = 7.5, 2% SDS, 0.5 mM EDTA, 350 mM 2-mercaptoethanol).

Tryptic digestion of the samples was performed using S-Trap Micro Spin Columns (ProtiFi, USA) according to the manufacturer’s instructions. Briefly, samples were acidified by adding phosphoric acid to a final concentration of 1.2%, 7 volumes of binding buffer (90% MeOH, 100 mM TEAB, pH = 7.1) was added to the samples, which were then transferred to the S-Traps, and spun at 4,000 x g for 30 seconds. The trapped proteins were washed three times with the binding buffer. Protein digestion was performed by adding 2 µg trypsin (Pierce, Thermo Scientific 90057) in 125 µL of 50 mM TEAB and incubating for 2 hours at 47 °C. Peptides were eluted with 0.2% FA and 0.2% FA/50 % ACN and dried down using vacuum centrifugation. Next, samples were resuspended in 40 μL of 0.1% FA and 10 μL was loaded onto disposable LC trap tips (Evotip).

NanoLC-MS/MS analysis was performed on an Evosep One UPLC system (Evosep) coupled to an Orbitrap Fusion Lumos Tribrid mass spectrometer equipped with a FAIMS Pro ion mobility device (both from Thermo Fischer Scientific). Peptides were separated on an Endurance 30SPD (Evosep) C18 analytical column (1.9 µm, 150 µm i.d. × 15 cm) using the 44-min “30 sample per day” non-linear gradient of Evosep One. Solvent “A” consisted of 0.1% FA, while solvent “B” contained 0.1% FA in ACN. The column temperature was 30 °C. A DDA method was applied, where MS1 full scans were acquired with a resolution of 60,000 (@ 200 m/z), applying custom normalized target AGC values of 50%, and a maximum injection time of 250 milliseconds, using a mass range of m/z 300-1500. The most intense multiply charged ions (z = 2-5) were fragmented using HCD activation applying stepped collision energies of NCE = 32 and 35. MS2 scans were acquired in the linear ion trap with a rapid scan rate, applying standard AGC target and a maximum IT of 100 milliseconds. The ion selection threshold was set to 1E+04 and the dynamic exclusion to 60 seconds. Data were acquired in 1.5 second cycles applying alternating FAIMS CV voltages of -70 or -50V.

Protein identification and label-free quantification was performed using Proteome Discoverer v3.0 SP1 (Thermo Fisher Scientific) with SEQUEST HT as the search engine and a human protein database downloaded from SwissProt on 2023-09-13 concatenated with a „common contaminants” database. Trypsin was specified as the protease, allowing for up to one missed cleavage. The mass tolerance was set to 5 ppm for MS1 and 0.6 Da for MS2. Methylthio modification of cysteine was set as a static modification, while methionine oxidation, pyroglutamine formation of peptide N-terminal glutamine, and protein N-terminal acetylation were included as dynamic modifications. Peptides and corresponding proteins were identified at a 1% false discovery rate (FDR). High-confident peptide identifications of Sequest XCorr Score>=1 were used to calculate abundance ratios across the sample groups using the „unique+razor” approach. Peptide abundances were normalized using the total peptide amount approach. Protein ratio calculation was pairwise ratio-based, and the hypothesis test was a background-based t-test. Expression levels of |log2FC|>=1 with adjusted p value=<0.05 were accepted as significant.

Protein abundance data from the co-immunoprecipitation experiment was processed using R v4.3.1. All proteins identified with “High” confidence, i.e., with an FDR < 0.05 and not classified as contaminants were filtered. Further, given that the iN samples were not pure cultures and still retained a large proportion of fibroblasts, all proteins identified as differentially abundant in the anti-IgG Immunoprecipitation of Ctrl fibroblasts when compared to that of Ctrl-iNs (Supplementary Table 14), were further excluded as background. Additionally, only proteins quantified in at least 3 samples belonging to one group and with an adjusted p-value < 0.05 in the pairwise ratio test were considered significant.

### Statistical analysis

In the iN morphology and MXRA8 spot analyses, each dot represents an HD or Ctrl donor cell line. Each of these values are an average of several individual cells from at least two individual wells (technical duplicates). The average relative value of a donor cell line was defined as the average value of all wells per line divided by the average value of all control wells. The acquisition and analysis of all wells was performed with identical settings of the HCA microscopy. All represented neuronal profiling values of a donor cell line (cell body area, neurite count, width, length, area and branchpoint count) are average relative values of several individual cells from at least two individual wells. The average relative dot number and area values of a donor cell line in autophagy analysis was defined as the average dot number or area of all wells per donor cell line divided by the average dot number and area of all control wells. The acquisition and analysis of all wells was performed with identical settings of the HCA microscopy.

To test differences between the two groups two-tailed unpaired or paired t-tests were used. For comparison of more than two groups, one-way ANOVA or non-parametric Kruskal-Wallis tests were used according to testing of distribution of the data by Saphiro-Wilk or D’Agostino-Pearson omnibus normality test. Data are presented as dots and means with error bars representing standard error of the mean (SEM) as specified in each figure legends.

## RESULTS

### HD-iNs display a distinct phosphoproteomic profile

We performed mass spectrometry-based phosphoproteomics (P-MS) on iNs derived from fibroblasts of seven individuals diagnosed with HD and seven age-matched controls (Fig. 1A and Table 1). These iNs, previously characterized in our 2022 study (Pircs et al. [22]), were generated using a well-established protocol [21, 26, 36, 37]. Our earlier findings showed that iNs from HD donors exhibit age-related epigenetic signatures, distinct proteomic profiles, and impaired autophagy [22].

**Figure 1.**
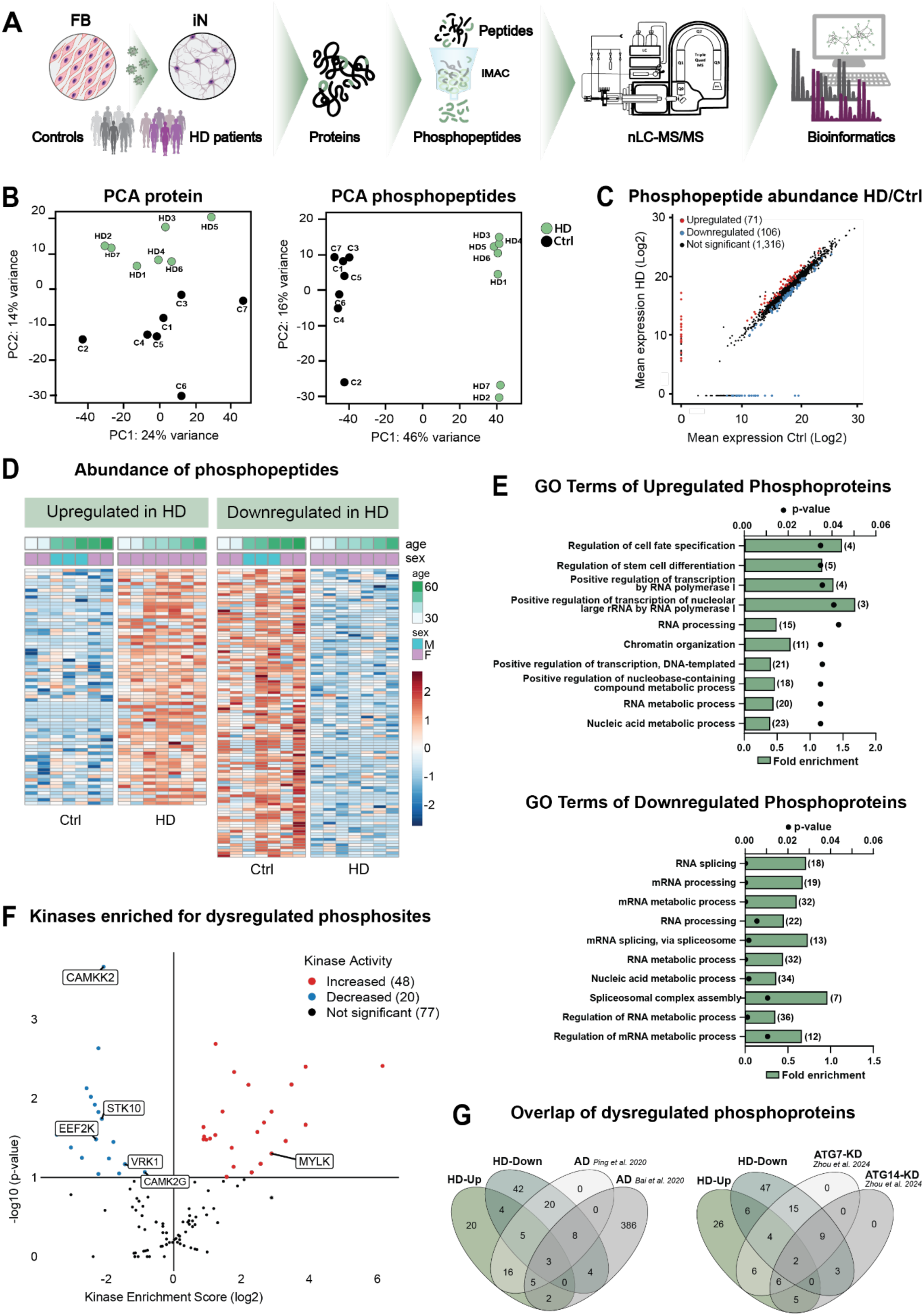
HD-iNs show a distinct phosphoproteomic profile. (A) Schematic overview of the experimental procedure. (B) PCA plots of nanoLC/MS and Phospho-nanoLC/MS data of iN samples grouped by Control (Ctrl) (n = 7) and HD (n = 7). (C) Scatter plot displaying log2 means of abundance of phosphopeptides in iNs from Ctrl (x-axis) and HD (y-axis) samples. Phosphopeptides significantly upregulated in HD-iNs when compared to Ctrl-iNs are represented by red dots, while those that are significantly downregulated are represented by blue dots, and the non-significant ones by black dots (Shapiro-wilk test p > 0.05; unpaired t-test p < 0.05). (D) Heatmaps showing the abundance of upregulated phosphopeptides (left) and downregulated phosphopeptides (right) in each sample. (E) Gene Ontology over-representation testing (using STRING-db v12.0) of genes associated with the upregulated phosphopeptides (top) and downregulated phosphopeptides (bottom). The top ten most significant terms are shown. Bar plots represent fold enrichment (strength), dots represent Benjamini-Hochberg false discovery rates (n = 7 control and 7 HD-iN lines, FDR<0.05). (F) Volcano plot showing the predicted kinases enriched for the phosphosites associated with the differentially abundant phosphopeptides. Kinases predicted to have an increased activity are shown in red, while those with decreased activity are shown in blue. Among other factors, significance was measured at p<0.05. (G) Venn diagrams showing the number of intersections of proteins associated with upregulated and downregulated phosphopeptides in the HD-iNs with those in (left) autophagy disruption experiments in hiPSC-iNs, and (right) studies of Alzheimer’s disease in post mortem frontal cortex samples.

We identified a total of 5,658 phosphopeptides and 4,866 assigned phosphosites. The characteristics of these phosphopeptides, including their detection frequency across samples and distribution of phosphosites per peptide, are summarized in Supplementary Fig. 1A and B. After filtering for phosphopeptides present in at least four samples and correcting for the intensities based on matching proteomic data (Methods), 4,183 phosphopeptides corresponding to 1,409 unique phosphoproteins were retained for downstream analysis (Supplementary Table 1).

Principal component analysis (PCA) revealed greater separation between HD-iNs and Ctrl-iNs in the phosphoproteomic data compared to the proteomic profiles of the same samples presented in Pircs et al 2022 [22], suggesting more pronounced post-translational differences in HD (Fig. 1B). Differential abundance analysis identified 177 significantly altered phosphopeptides corresponding to 129 proteins. Among these, 71 phosphopeptides (from 55 proteins) were upregulated, and 106 phosphopeptides (from 86 proteins) were downregulated in HD-iNs compared to Ctrl-iNs (Fig. 1C and D; Supplementary Table 2 and 3). Notably, 12 phosphoproteins had both upregulated and downregulated phosphopeptides. While, we detected phosphopeptides from HTT (specifically Ser417, Ser419, and Ser432), which have previously been linked to disease pathology [38–40], their abundance was not significantly different between HD-iNs and Ctrl-iNs.

Interestingly, comparison with proteomic and transcriptomic datasets from Pircs et al [22], revealed limited overlap with the phosphoproteomic changes (Supplementary Fig. 1C-F). Out of 129 genes corresponding to differentially abundant phosphopeptides, only a small subset showed altered expression at other molecular levels. Eight genes with downregulated phosphopeptides were upregulated at the proteomic level in HD-iNs, namely, *AKAP8*, *ARHGEF6*, *FTO*, *MXRA8, NFXL1*, *SF3A3*, *TMEM214* and *ZNF22* (Supplementary Fig. 1C and D). Among these, several - *ARHGEF6*, *AKAP8*, *SF3A3*, *NFXL1*, *ZNF22*, and *FTO* - are involved in transcriptional regulation, RNA processing, or neuronal signaling [41–50]. This suggests dysregulation of nuclear and synaptic pathways at the post-translational level, with the increased protein abundance potentially reflecting a compensatory response to impaired post-translational control mechanisms in HD. In addition, five genes, namely, *AKAP8*, *NFXL1* and *STRIP1* with downregulated phosphopeptides, and *TCOF1* and *ST5*, with upregulated phosphopeptides, were upregulated at the transcriptomic level (Supplementary Fig. 1E and F). Notably, only *AKAP8* and *NFXL1* were dysregulated across all three expression levels: transcriptomic, proteomic, and phosphoproteomic (Supplementary Fig. 1D and F). *AKAP8* and *NFXL1* are both involved in RNA processing and transcriptional regulation [45, 46, 48], suggesting they may play key roles in mediating HD-related changes at the post-transcriptional level.

Furthermore, the presence of Serine/Arginine Repetitive Matrix proteins (SRRM1 and SRRM2) in human HD-iNs, combined with the identification of phosphoproteomic changes previously reported in their mouse homologues (SRRM1, SRRM2) as well as in CLIP2, MAP1B, PI4KB, and Synaptopodin (SYNPO), suggests conserved disruption of splicing pathways [51, 52]. Together, these observations provide additional support that HD involves pronounced post-transcriptional and post-translational alterations, consistent with known splicing and RNA processing disruptions in HD [53–57].

### Functional annotation reveals dysregulated kinase activity and pathways

To better understand the functional consequences of the phosphoproteomic changes, we performed a functional enrichment analysis on the proteins associated with the dysregulated phosphopeptides, using STRING-db. Gene Ontology (GO) terms for the upregulated phosphopeptides were significantly enriched for biological processes such as cell fate specification and transcriptional regulation, while those for the downregulated phosphopeptides were primarily associated with RNA splicing (Fig. 1E and Supplementary Tables 4 and 5). Reactome pathway terms showed an enrichment for autophagy-related pathways, including PTEN regulation and PIP3-AKT signaling, for the upregulated phosphopeptides (Supplementary Fig. 1 G and H, Supplementary Tables 6 and 7), suggesting a potential post-translational contribution to autophagy dysregulation in HD-iNs.

We next performed Kinase Enrichment Analysis using the Kinase Library’s PSEA module on PhosphoSitePlus [30–32] to identify potential kinases responsible for the observed phosphopeptide changes. Among the predicted kinases overlapping with those previously quantified in the proteomic dataset [22], we identified 48 with increased activity and 20 with decreased activity in the HD-iNs (Fig. 1F and Supplementary Table 8). The kinases with increased activity predominantly belonged to the CAMK and AGC families, while those with decreased activity were largely from the STE family (Supplementary Fig. 2A).

Of the kinases previously found to be significantly differentially abundant at the proteomic level in the HD-iNs, only 4 showed both predicted kinase activity and changes in protein abundance in the same direction. These included STK10, CAMK2G, CAMKK2, and EEF2K, all with decreased predicted activity and downregulation in the HD-iNs (Fig. 1F, Supplementary Fig. 2B, Supplementary Table 8). The remaining predicted kinases that were differentially abundant, - VRK1 and MYLK - did not show consistent trends between predicted activity and protein levels (Supplementary Table 8).

Together, these findings highlight widespread alterations in signaling and phosphorylation regulation in HD-iNs, driven in part by dysregulated kinase activity across multiple functional families. Notably, many of the kinases with altered activity are linked to autophagy-related pathways, further supporting the concept that post-translational dysregulation of autophagy is a central feature of HD-iNs.

### Shared phosphoproteomic signatures with autophagy disruption and neurodegeneration

To determine whether the phosphoproteomic alterations observed in HD-iNs are shared with other models of neurodegeneration, we compared our dataset with phosphoproteomic profiles from three previously published sources.

First, we found substantial overlap with phosphoproteomic changes observed in autophagy-disrupted human iPSC-derived neurons with CRISPRi-mediated knockdown of *ATG7* and *ATG14*, as described by Zhou et al., 2024 [58] (Supplementary Table 9). Specifically, 27 phosphoproteins with downregulated phosphopeptides and 17 with upregulated phosphopeptides in HD-iNs were also dysregulated in these autophagy-deficient models (Fig. 1G, Supplementary Fig. 2C and Supplementary Table 9). This suggests that phosphoproteins altered in HD-iNs may contribute to, or be influenced by, autophagy dysfunction.

Second, we observed that 32 phosphoproteins with downregulated phosphopeptides and 23 with upregulated phosphopeptides in HD-iNs overlapped with phosphoproteomic alterations in post mortem frontal cortices of Alzheimer’s disease (AD) patients [59, 60] (Fig. 1G, Supplementary Fig. 2C and Supplementary Table 10). This suggests that similar signaling pathways may be disrupted across different age-related neurodegenerative diseases, highlighting the broader importance of post-translational dysregulation in these conditions.

### ON-OFF and OFF-ON phosphoproteins reveal binary phospho-switches in HD-iNs

We identified a subset of 27 downregulated phosphopeptides, corresponding to 26 phosphoproteins, that were entirely absent in HD-iNs but present in at least four Ctrl-iNs (Fig. 2A and Supplementary Table 11). We refer to these as ON-OFF phosphopeptides/proteins.

**Figure 2.**
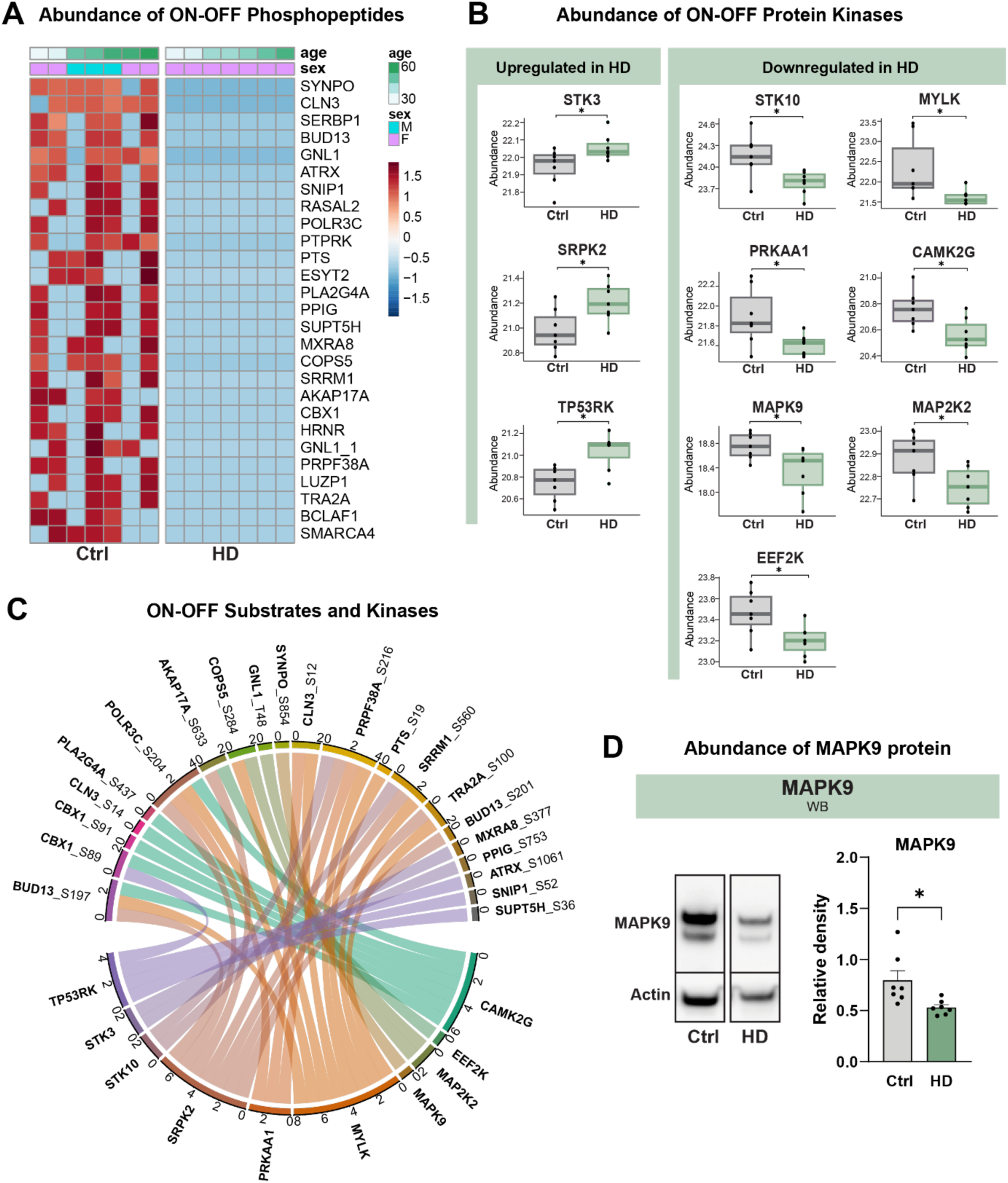
ON-OFF proteins and their predicted kinases. (A) Heatmap showing the abundance of the ON-OFF phosphopeptides in each sample. (B) Box plots showing the abundance of the kinases, predicted to phosphorylate ON-OFF phosphosites, that are upregulated in HD-iNs when compared to Ctrl-iNs (left), or downregulated in HD-iNs (right). (n = 7 control and 7 HD iN donor cell lines) (C) Chord diagram summarizing the interactions between the predicted kinases and the corresponding ON-OFF phosphosites. The scales represent the number of connections. (D) Validation of MAPK9 protein abundance by WB of 7 Ctrl and 7 HD-iNs. (B, D) Each dot represents one Ctrl or one HD adult human donor cell line from the converted iNs. For statistical analysis two tailed t-tests were used in all cases. * p < 0.05 Data is shown as mean ± SEM in D.

Except for MXRA8, which was upregulated at the proteomic level (Fig. 3B), none of the ON-OFF proteins showed significant changes at either the transcriptomic or proteomic levels (Supplementary Fig. 3A and B, Supplementary Table 11). Notably, two ON-OFF phosphoproteins - SYNPO and SRRM1 - have previously been reported to be dysregulated in the striatal and hippocampal phosphoproteomes of HD mouse models [12, 13]. Furthermore, SRRM1, along with nine additional ON-OFF phosphoproteins (CLN3, SERBP1, SNIP1, ESYT2, PPIG, SUPT5H, PRPF38A, BCLAF1, and SMARCA4), were also found to be dysregulated in autophagy-deficient iPSC-derived neurons from *ATG7* and *ATG14* knockdown experiments [58] (Supplementary Fig. 3F and Supplementary Table 9). Additionally, SYNPO, SRRM1, BCL2 Associated Transcription Factor 1 (BCLAF1) and SWI/SNF Related BAF Chromatin Remodeling Complex Subunit ATPase 4 (SMARCA4) were also identified to be dysregulated in the post mortem frontal cortex phosphoproteomes of AD patients [59, 60] (Supplementary Fig. 3F and Supplementary Table 10). Among these, SRRM1 is of particular interest, as it is a core component of the splicing machinery. Its phospho-dysregulation links RNA processing to both autophagy impairment and age-related neurodegeneration and highlights the intersection of post-transcriptional regulation with disease-relevant cellular pathways [61, 62].

**Figure 3.**
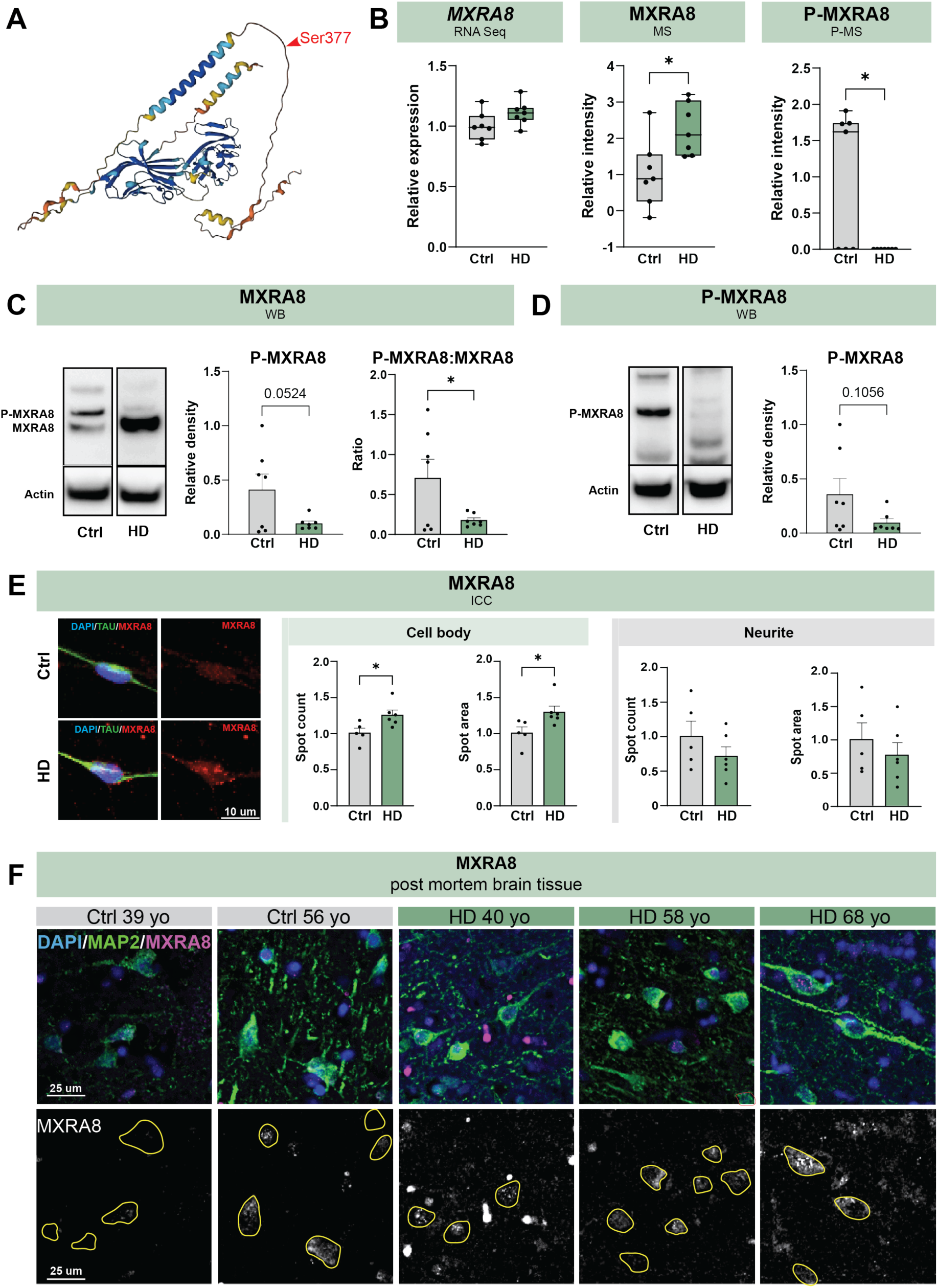
HD-iNs show dysfunction of MXRA8 protein. (A) 3D structural image of the MXRA8 protein from Alphafold. Red arrow points to the specific phosphosite Ser377 which is completely diminished in HD-iNs. (B) RNA expression, protein abundance and phospho-protein abundance of MXRA8 in 7 Ctrl- and 7 HD-iNs from RNA-seq, MS and P-MS analysis, respectively. (C) WB validation of MXRA8 protein abundance of n=13 of N=7 Ctrl- (two biological replicates per each line, except for C2) and n=14 of N=7 HD-iNs (two biological replicates per each line) using native anti-MXRA8 (“N” represents the number of donor cell lines, “n” represents the total number of biological replicates). Blot sections show one representative Ctrl- and HD-iN samples. Column diagrams show the relative density of P-MXRA8 and a ratio of P-MXRA8:MXRA8 as analyzed on the native MXRA8-labelled blots. (D) WB validation of P-MXRA8 protein abundance of n=14 of N=7 Ctrl and n=14 of N=7 HD iNs using custom designed phospho-Ser377-MXRA8 specific antibody. Blot sections show one representative Ctrl- and HD-iN samples. Column diagrams show the relative density of P-MXRA8. (E) Validation of MXRA8 protein abundance and localization by HCA in TAU counterstained 7 Ctrl- and 7 HD-iN samples. Bars show the spot count and area in the cell body (left) and in the neurites (right) of Ctrl- and HD-iNs. Representative images show one Ctrl- and one HD-iN sample stained with DAPI, TAU and MXRA8. (F) Representative images of IHC validation of MXRA8 protein expression in post mortem human cortical brain samples from 2 Ctrl and 3 HD individuals counterstained with MAP2 and DAPI. Yellow outlines mark neuronal cell bodies based on MAP2 staining. Scale bar 25 μm. (B-E) Each dot represents one Ctrl- or one HD-adult human cell line from the converted iNs, the columns of the bars show the mean value of the individual values, the error bar indicates SEM. For statistical analysis two tailed t-tests were used, * p<0.05.

We additionally identified a subset of 19 OFF-ON phosphopeptides, corresponding to 17 phosphoproteins, that were completely absent in Ctrl-iNs but expressed in at least four HD-iN samples (Supplementary Fig. 3C and Supplementary Table 12). None of the genes corresponding to these phosphopeptides were dysregulated at other levels of expression in the HD-iNs, except for Treacle Ribosome Biogenesis Factor 1 (TCOF1), which was upregulated at the transcriptomic level (Supplementary Fig. 3D and E). Among these, CLIP2 (CAP-Gly domain-containing linker protein 2) was previously reported as dysregulated in the phosphoproteome of HD mouse models [12, 13]. In addition, TCOF1 and CLIP2, - as well as IWS1, NDRG2, NCL, and UBE2O - were also found to be altered in the phosphoproteomes of autophagy-disrupted iPSC-derived neurons and post mortem frontal cortices from AD patients (Supplementary Fig. 3F, Supplementary Table 9 and 10) [58–60].

Together, the ON-OFF and OFF-ON phosphoproteins represent strong, binary phospho-switches in HD-iNs, marked by either a complete presence or complete absence of phosphorylation. The fact that many of these proteins are also found in HD models, autophagy-impaired neurons, and AD brain tissue suggests they may play important roles in regulating cell signaling and stress responses in neurodegenerative conditions.

### Dysregulated ON-OFF kinases link phospho-switches to autophagy and neurodegenerative signaling

Using the kinase prediction tool from the PhosphoSitePlus database [30–32], we identified potential upstream kinases for the ON-OFF phosphosites (Supplementary Table 13). Overall, we considered kinases that were expressed in the global proteomic dataset and had a site percentile greater than 90, as potential kinases. Further, among those, 10 kinases were differentially abundant in the global proteomic data, where STK3, SRPK2 and TP53RK, were upregulated in the HD-iNs, while STK10, PRKAA1, MYLK, MAPK9, CAMK2G, MAP2K2 and EEF2K were downregulated (Fig. 2B). Several of these differentially abundant kinases were linked to multiple ON-OFF phosphosites, highlighting their potential regulatory significance.

Notably, PRKAA1 (AMPKα1), a central kinase in the CAMKK-AMPK autophagy pathway, along with MYLK and CAMK2G, was predicted to phosphorylate CLN3, a core autophagy-associated protein [63], at serine 12 (S12), and CAMK2G at S14 (Fig. 2C). These observations suggest that reduced levels of PRKAA1, MYLK, or CAMK2G may underlie the lack of CLN3 phosphorylation in HD-iNs. Importantly, this links the dysregulation of autophagy-related signaling directly to the ON-OFF phosphoproteome.

While some of the ON-OFF phosphosites had multiple predicted kinases, two showed unique predicted kinase-substrate relationships: SYNPO-S854, phosphorylated by Mitogen-Activated Protein Kinase 9 (MAPK9), and AKAP17A-S633, by EEF2K (Fig. 2C), also known as JNK2 (c-Jun N-terminal kinase 2). MAPK9 is a stress-activated kinase with established roles in oxidative stress responses, apoptosis, and neurodegeneration [64, 65] [66]. WB analysis confirmed reduced MAPK9 protein levels in HD-iN compared to Ctrl-iNs, consistent with our proteomic results (Fig. 2D).

SYNPO, the substrate of MAPK9 at S854, is a cytoskeletal-associated protein highly enriched in neurons [67–71]. It plays a critical role in dendritic spine structure, actin organization, and synaptic plasticity – all processes vulnerable in neurodegenerative diseases [67, 72–74]. Its absence of phosphorylation in HD-iNs, paired with reduced MAPK9, may reflect impaired structural and signaling integrity in disease-relevant neuronal compartments.

In summary, kinase prediction analysis of ON-OFF phosphoproteins uncovered multiple potential regulators with strong connections to autophagy and neurodegenerative pathways. Dysregulation of key kinases such as MAPK9, PRKAA1, MYLK, and CAMK2G, and their links to substrates like CLN3 and SYNPO, support the hypothesis that loss of site-specific phosphorylation contributes to impaired signaling in HD-iNs, with broader implications for cellular stress response and synaptic function.

### Matrix-remodeling protein 8 shows distinct alterations in HD-iNs

We selected MXRA8 for further investigation, as it was the only ON-OFF phosphoprotein that also showed a significant change in overall protein abundance in HD-iNs (Fig. 3A and B). Although the phospho-Ser377 peptide of MXRA8 was completely absent in HD-iNs, the overall protein levels of MXRA8 were increased, as detected by conventional MS (Fig. 3B). The observed increased expression of the MXRA8 protein could potentially be a compensatory response to the loss of phosphorylated MXRA8. Given that the RNA sequencing data revealed no significant difference in *MXRA8* mRNA expression, the observed differences likely arise from post-transcriptional regulation (Fig. 3B). Notably, MXRA8 had only one phosphopeptide with a single phosphosite detected across the entire dataset, making it uniquely suited for targeted follow-up studies. We hypothesize that the combination of phosphorylation loss and protein-level upregulation may reflect a dysregulated state of MXRA8 in the HD-iNs, pointing to a possible role in disease-related mechanisms.

MXRA8 - also known as Limitrin, DICAM (dual immunoglobulin domain-containing cell adhesion molecule) or ASP3 (adipocyte specific protein 3) - is a transmembrane protein belonging to the immunoglobulin superfamily [23]. It is involved in various cellular processes, including mesenchymal cell differentiation, osteoclast development, angiogenesis, and viral entry [23, 75–77]. In the central nervous system, MXRA8 has been primarily described in glial cells, where it contributes to blood-brain barrier integrity and modulates astrocyte-mediated neuroinflammation [23, 78]. However, its role in neurons remains largely unknown. Our findings - showing a strong imbalance between MXRA8 phosphorylation and protein abundance in HD-iNs – suggest a previously unrecognized role for MXRA8 in neurons and suggest its possible involvement in neurodegenerative processes.

Therefore, we first validated the MS and P-MS data by measuring the abundance and ratio of MXRA8 and P-MXRA8 by WB using either a native anti-MXRA8 antibody (labelling both native and post-translationally modified forms of MXRA8) or our custom-designed phospho- Ser377-MXRA8 specific antibody (Fig. 3C and D, Supplementary Fig. 4A). Based on the evaluation of the native anti-MXRA8 labelled blots, HD-iNs showed a lower abundance of P-MXRA8 compared to Ctrl-iNs, the difference of which was close to significance (p = 0.0524). By contrast, the ratio of P-MXRA8:MXRA8 protein abundance showed a significant decrease in HD-iNs compared to non-diseased controls. The P-MXRA8 detection using the phospho-Ser377-MXRA8 specific antibody also confirmed a decreased abundance of P-MXRA8 in HD- iNs seen by the native anti-MXRA8 antibody (Fig. 3D). The lack of statistical significance for total MXRA8 in WB likely reflects the lower sensitivity of bulk WB compared to MS or ratio analyses for subtle changes.

Next, we performed ICC staining followed by high-content microscopy to visualize MXRA8, using the native anti-MXRA8 antibody and counterstaining with TAU (Fig. 3E, Supplementary Fig. 4B). MXRA8 staining appeared as punctate signals, primarily localized in the cell body and nuclei but was also present in the neurites of iNs (Fig. 3E, Supplementary Fig. 4B). In HD-iNs, the size and number of MXRA8-positive puncta were significantly increased in the cell bodies. In contrast, the neurites showed a trend toward fewer MXRA8 puncta, although this difference did not reach statistical significance (Fig. 3E). To further validate the neuronal relevance of these findings, we performed immunohistochemistry in post mortem cortical tissue from three advanced-stage (grade 4) HD patients and two age-matched controls, using MAP2 as a neuronal marker alongside MXRA8 (Fig. 3F). Similar to the HD-iNs, we observed a proportion of neuronal cell bodies with dense punctate MXRA8 staining in HD patient tissue.

Although the limited availability of tissue and small sample size prevented reliable quantification, the consistent presence of MXRA8 puncta in HD neurons strongly supports the notion that MXRA8 dysregulation observed in HD-iNs is also detectable in patients’ brain tissue.

These findings reveal a marked imbalance between MXRA8 phosphorylation and protein abundance in HD-iNs, accompanied by distinct subcellular redistribution and reflected by similar punctate MXRA8 accumulation in human HD cortical neurons, supporting a potential role for MXRA8 dysregulation in HD-related neuronal processes.

### MXRA8-associated interactome and kinases suggest a role in autophagy and neurovascular function

To explore the molecular context of MXRA8 and gain insight into its potential functions in neurons, we performed co-IP using the anti-MXRA8 antibody, followed by LC-MS/MS, on four control and four HD-iNs (Fig. 4A and Supplementary Table 15). Overall, we identified 1,232 proteins with high confidence (FDR < 0.05; Fig. 4B). From these, we shortlisted 20 candidate interactors based on significant abundance ratio, and reliable peptide-level quantification in at least three samples within one condition (Fig. 4B and Supplementary Tables 16). Of these, five proteins were robustly represented in ≥50% of samples in either group (Fig. 4B).

**Figure 4.**
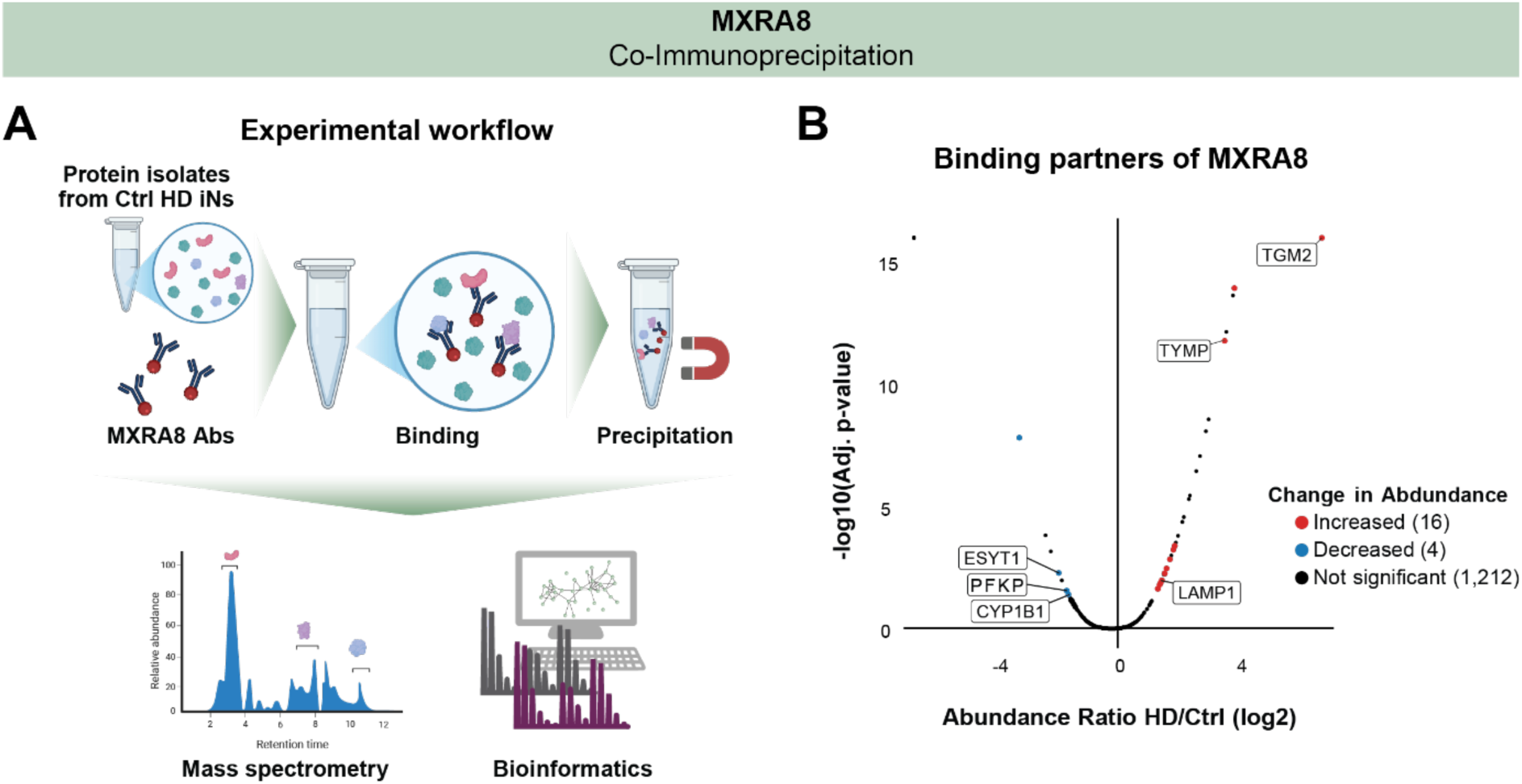
Co-immunoprecipitation identifies binding partners of MXRA8. (A) Schematic overview of the co-immunoprecipitation experimental procedure (B) Volcano plot of identified binding partners of MXRA8 of n=4 Ctrl- and n=4 HD-iN samples. On the volcano plot binding partners with increased abundance in HD-iNs are shown in red, while those with decreased abundance are shown in blue and the non-significant ones by black dots. Significance was measured based on an adjusted p-value < 0.05 among other parameters.

Among the candidates, Transglutaminase 2 (TGM2) and Lysosomal-associated membrane protein 1 (LAMP1) showed increased binding to MXRA8 in HD-iNs, while Cytochrome P450 Family 1 Subfamily B Member 1 (CYP1B1), Phosphofructokinase (PFKP), and Extended Synaptogamin 1 (ESYT1) were decreased compared to controls (Fig. 4B and Supplementary Table 16). TGM2 is known to modulate neurogenesis and synaptic pruning as well as mediating autophagosome-lysosome fusion, while LAMP1 is a key component of macroautophagy [79–84]. CYP1B1 is involved in blood-brain barrier integrity and inflammatory responses, PFKP is a critical glycolytic enzyme, and ESYT1 regulates calcium transport at membrane contact sites [85–88]. These changes in binding profiles support a broader role for MXRA8 in maintaining neuronal homeostasis and metabolic balance.

Due to the lack of experimental studies on MXRA8 phosphosites, the kinases regulating its phosphorylation remain unidentified. Among the list of kinases of MXRA8-pS377 that we had predicted earlier with a site percentile greater than 90 and expressed in the global proteomic data (Supplementary Table 13), AAK1, BMP2K, GAK, PRKCD, PRKCA, IRAK4, DSTYK, PRKCE, STK26 and EIF2AK2 had the highest site scores (Supplementary Fig. 4C). The site score reflects the similarity between the phosphosite and a given kinase’s preferred consensus motif, making it an estimate of how likely the kinase is to phosphorylate the phosphosite. Further experiments would need to be performed to further validate the role of these kinases in the phosphorylation of MXRA8-pS377.

Together, these findings suggest that altered kinase activity and changes in MXRA8 protein-protein interactions may influence its phosphorylation state and function in HD-iNs. The association of MXRA8 with proteins involved in metabolism, calcium signaling, and the lysosomal/autophagy axis, along with potential regulation by autophagy-linked kinases, points to a broader role in neurovascular integrity and stress response pathways relevant to neurodegeneration.

## DISCUSSION

Although the causative genetic mutation was recognized over thirty years ago, the exact pathogenesis of this condition remains unknown [2]. Epigenetic hallmarks and neuronal age are both key to HD pathology as symptoms occur only after a certain age despite the mutation being inherited [19]. In this study we used our previously published HD-iN proteomic and transcriptomic dataset and expanded it with P-MS measurements to investigate the phosphoproteomic profiles of patient-derived directly reprogrammed HD-iNs. Importantly, phosphoproteomics provides a dynamic view of protein regulation that is not captured by steady-state transcriptomic or proteomic analyses, allowing us to study key signaling disruptions that may precede overt disease phenotypes. HD-iNs not only retains the patients genetic mutation but also recapitulates the epigenetic age of the donor [19, 22, 36]. We previously found that HD-iNs: *i)* mainly differ on a proteomic, but not so much on a transcriptomic level, *ii)* show a less elaborate neuronal morphology, *iii)* show an accelerated epigenetic aging based on DNA methylation array, and *iv)* have clear autophagy dysfunction [22].

While growing knowledge is available about the DNA methylation, transcriptomic, and proteomic profiles of HD, much less is known about the contribution of post-translational phosphorylation dysregulation to the disease mechanism [89, 90]. Previously published studies focusing on protein phosphorylation in HD mainly studied the importance of specific protein phosphorylation sites on HTT, TAU or histone proteins [9–11, 14, 15, 91, 92]. Here we performed an untargeted phosphoproteomic profiling approach using seven HD and seven Ctrl- iNs. This comprehensive, unbiased analysis of the phosphorylation network enabled us to identify new key proteins and dysregulated pathways, especially when combined with our previously published transcriptomic and proteomic datasets from the same patient-derived iNs and controls.

So far, such analyses have only been done in mouse models of HD. The first study published by Beaumont et al in 2016 found no significant change in the phosphoproteome of the striatum of Q175 HD mice [93]. In 2022 Mees et al. performed P-MS analysis of different brain regions of R6/1 transgenic mice at premanifest and manifest stages [12, 13]. They reported strong alterations in phosphoprotein levels in premanifest mice, particularly in the striatum and the cortex, but with less differences at manifest stages [12, 13].

The phosphoproteomic analysis presented in our study is the first phosphoproteome-wide investigation of patient-derived human samples. P-MS data showed a clear distinction between HD-iNs and Ctrl-iNs at the phosphoproteomic level. Interestingly, we only found four overlapping phosphoproteins between our HD-iN dataset and the HD mouse phosphoproteome from Mees et al. (MAP1B, PI4KB, SRRM1, and SRRM2) [12, 13]. While inconsistencies between the nomenclature of mice and human proteins may explain part of this discrepancy, the limited overlap strongly highlights the importance of studying HD in human derived models. Mees et al. also reported that pathways such as neurite outgrowth, ubiquitination, tight junction assembly, and endocytosis were upregulated, while mRNA translation, ion homeostasis, and amino acid transport were downregulated. Cytoskeleton organization and phosphatase activity were mixed across conditions [13].

STRING-db analysis of our HD-iN P-MS data showed a downregulation of RNA splicing pathways. Out of the 18 significantly downregulated proteins related to RNA splicing, 8 were identified as ON-OFF proteins (BUD13, SNIP1, PRPF38A, TRA2A, SRRM1, PPIG, AKAP17A and BCLAF1). Previously, we found RNA splicing to be significantly upregulated on a protein level (MS) [22]. One overlapping protein, SF3A3 (Splicing Factor 3a subunit 3) is present with significantly upregulated abundance but significantly decreased phosphorylation in HD-iNs, suggesting possible functional disruption. At the transcriptomic level, RNA splicing-related processes were found to be significantly up and downregulated, highlighting the complexity of post-transcriptional RNA regulation in HD-iNs [22]. These findings align with the current literature that implicate RNA splicing dysregulation as a key pathogenic mechanism in HD [53–56].

To contextualize our findings beyond HD models, we compared our dataset with other phosphoproteomic studies in human neurodegeneration, including AD and models of autophagy dysfunction. Since large-scale phosphoproteomic datasets for neurodegenerative conditions remain limited, we focused our comparisons on three high-confidence datasets: iPSC-derived neurons with CRISPRi-mediated knockdown of autophagy genes (ATG7 and ATG14) [58], and post mortem brain tissue from AD patients [58–60]. We found significant overlap in phosphoproteomic alterations between HD-iNs and both autophagy-impaired neurons and AD brains. Specifically, 44 phosphoproteins dysregulated in HD-iNs were also affected in the autophagy-deficient neurons, and 55 overlapped with those altered in AD brains. Notably, 22 phosphoproteins were altered across all three datasets, suggesting they may act as shared molecular mediators in age- and stress-related neurodegeneration.

Notably, MAP1B, SRRM1 and SRRM2 were significantly differently phosphorylated in the mouse HD brain as well [12, 13]. A subset of nine phosphoproteins showed conserved changes at the same phosphosite and in the same direction across these datasets, supporting the existence of conserved post-translational regulatory mechanisms. We also identified a class of ON-OFF phosphoproteins - those completely unphosphorylated in HD-iNs yet present in controls. Among these, several proteins (e.g., SRRM1, SYNPO, BCLAF1) were also ON-OFF in both autophagy-deficient neurons and AD brains, indicating that loss of site-specific phosphorylation may represent a common binary switch mechanism in neurodegenerative pathways.

Strikingly, only one of the ON-OFF proteins, MXRA8 had one single non-phosphorylated phosphosite - at Serine-377 - while having a significantly higher abundance in HD-iNs. Presumably, the diminished function of MXRA8 caused by interrupted phosphorylation resulted in a compensatory increase in protein production by the cells. Importantly, we also verified MXRA8 expression in neurons of post mortem HD cortical tissue, where we observed a trend toward increased punctate accumulation in neuronal cell bodies, consistent with our HD-iN findings. MXRA8 is a transmembrane protein of the immunoglobulin superfamily, previously characterized mainly in glial and vascular contexts, but not in neurons. Here, we demonstrate for the first time its altered phosphorylation and abundance in human neurons, implicating MXRA8 dysregulation in neurodegenerative processes [23, 94].

To explore the functional implications of MXRA8 dysregulation, we performed co-immunoprecipitation coupled with LC-MS/MS, identifying a selective shift in MXRA8’s binding partners in HD-iNs. Proteins involved in lysosomal function (LAMP1) and stress homeostasis (TGM2) showed increased interaction, while partners related to glycolysis (PFKP) and calcium transport (ESYT1) displayed reduced associations in HD-iNs. These context-specific interaction shifts suggest that MXRA8 participates in metabolic and stress-response pathways disrupted in HD.

Further, we predicted potential kinases regulating MXRA8 phosphorylation. Of particular interest were AAK1, BMP2K, GAK, PRKCD, PRKCA, IRAK4, DSTYK, PRKCE, STK26 and EIF2AK2, all previously implicated in autophagy regulation [95–104]. MXRA8 has no experimentally determined kinases, due to a lack of studies on MXRA8 phosphosites and the overall small fraction of known kinase-substrate relationships [105–107]. This limits our ability to identify potential kinases for MXRA8 based on established kinase prediction tools. Additional experimental studies that evaluate the potential of the shortlisted kinases in phosphorylating MXRA8 would therefore be required for further verification.

Taken together, our findings identify MXRA8 as a novel, neuronally relevant protein whose dysregulated phosphorylation and protein-protein interactions may contribute to autophagy impairment and neurovascular dysfunction in HD [108]. The loss of phosphorylation in combination with increased protein abundance, disrupted interactome, and reduced autophagy-linked kinase activity suggests a broader role for MXRA8 in neurodegenerative stress responses.

In summary, this study presents the first phosphoproteome-wide analysis of human HD neurons, using directly reprogrammed iNs from genetically and epigenetically characterized patient donors. By integrating phosphoproteomic, proteomic, and transcriptomic data from the same individuals, we demonstrate the critical role of PTMs - especially phosphorylation - in HD pathogenesis. The divergence between phosphorylation status and gene or protein expression in many key regulators, including MXRA8, highlights that PTMs provide distinct and functionally relevant disease signals that are not captured by transcriptomic or steady-state proteomic profiling alone. Our findings not only reveal previously unrecognized HD-associated phospho-switches and disrupted signaling pathways, particularly in autophagy and stress response networks, but also identify novel candidates such as MXRA8 for mechanistic follow-up and therapeutic targeting. This multi-omics approach establishes a new platform for identifying clinically relevant biomarkers and druggable targets in human neurodegeneration.

## DATA AND CODE AVAILABILITY

All data needed to evaluate the conclusions in the paper are present in the paper and/or the Supplementary Materials section.

The mass spectrometry phosphoproteomics data have been deposited to the ProteomeXchange Consortium via the PRIDE [109] partner repository. The mass spectrometry dataset of the co-IP experiment was uploaded to the Proteomics Identifications Database (PRIDE, https://www.ebi.ac.uk/pride/login). The code used for the data analysis and for generation of plots can be found at https://github.com/pircs-lab/iN_HD_PMS_analysis_2025.

## ACKNOWLEDGEMENTS

We are thankful to Sara Bermudez, Guilia Saredi, Johan Jakobsson and to all members of the HCEMM-SU Neurobiology and neurodegenerative diseases research group. We also thank Pálma Anna Zsolnai for her valuable help with lentiviral vector production at the Semmelweis Viral Vector Core.

## AUTHOR CONTRIBUTIONS

R.A.B. provided the human fibroblasts and post mortem material used in the study. J.G. carried out the mass spectrometry-based phosphoproteomic experiments under the supervision of M.R. and G.M.V. L.D. and C.M. designed and visualized the figures. C.M. designed the computational framework. C.M. and D.J. analysed the data under the supervision of K.P. L.D. designed, supervised and performed the experimental work together with Á.V., Á.S., A.A.A., and R.Z. G.R. designed, supervised and performed co-IP experiments together with M.C. Z.D. supervised, performed together with Á.P. and analysed together with C.M. the MS experiments using the co-IP samples. A.S.P. performed experiments using post mortem material under the supervision of J.D.O. L.D. and C.M. wrote the manuscript with support from K.P. and with input from all authors. K.P. and L.D. conceived the study and along with C.M. were in charge of overall direction and planning. K.P. and L.D. supervised the project. All authors provided critical feedback and helped shape the research, analysis and manuscript.

## ETHICS STATEMENT

Experiments with human fibroblast samples were conducted under ethical approval IV/2625-1/2021/EKU, and those involving human post mortem material under approval CERC-2025-7229 (University of Montreal).

## SUPPLEMENTARY INFORMATION

### Supplementary Data

Following data are provided in a separate .zip file:

**Supplementary Table 1:** Phosphopeptides identified across the HD-iNs and the Ctrl-iNs

**Supplementary Table 2:** Differential abundance analysis results: Phosphopeptides upregulated in HD-iNs vs Ctrl-iNs.

**Supplementary Table 3:** Differential abundance analysis results: Phosphopeptides downregulated in HD-iNs vs Ctrl-iNs.

**Supplementary Table 4:** Gene ontology terms enriched for phosphoproteins associated with phosphopeptides upregulated in HD-iNs vs Ctrl-iNs.

**Supplementary Table 5:** Gene ontology terms enriched for phosphoproteins associated with phosphopeptides downregulated in HD-iNs vs Ctrl-iNs.

**Supplementary Table 6:** Reactome pathways enriched for phosphoproteins associated with phosphopeptides upregulated in HD-iNs vs Ctrl-iNs.

**Supplementary Table 7:** Reactome pathways enriched for phosphoproteins associated with phosphopeptides downregulated in HD-iNs vs Ctrl-iNs.

**Supplementary Table 8:** Kinase enrichment analysis results for dysregulated phosphopeptides in HD-iNs vs Ctrl-iNs

**Supplementary Table 9:** List of phosphoproteins dysregulated in the HD-iNs overlapping with phosphoproteins dysregulated in ATG7- and ATG-14 knock-down studies from Zhou et al. 2024

**Supplementary Table 10:** List of phosphoproteins dysregulated in the HD-iNs overlapping with phosphoproteins dysregulated in post mortem Alzheimer’s disease brain samples from Bai et al. 2020 and Ping et al. 2020

**Supplementary Table 11:** List of ON-OFF phosphopeptides

**Supplementary Table 12:** List of OFF-ON phosphopeptides

**Supplementary Table 13:** Kinase prediction results for ON-OFF phosphoproteins

**Supplementary Table 14:** List of binding partners of MXRA8 in fibroblast samples identified through anti-MXRA8 co-Immunoprecipitation coupled with LC-MS/MS (pilot experiment)

**Supplementary Table 15:** List of binding partners of MXRA8 in induced neurons identified through anti-MXRA8 co-Immunoprecipitation coupled with LC-MS/MS

**Supplementary Table 16:** Filtered list of binding partners of MXRA8 in induced neurons identified through anti-MXRA8 co-Immunoprecipitation coupled with LC-MS/MS

## Supplementary Materials

**Supplementary Figure 1.**
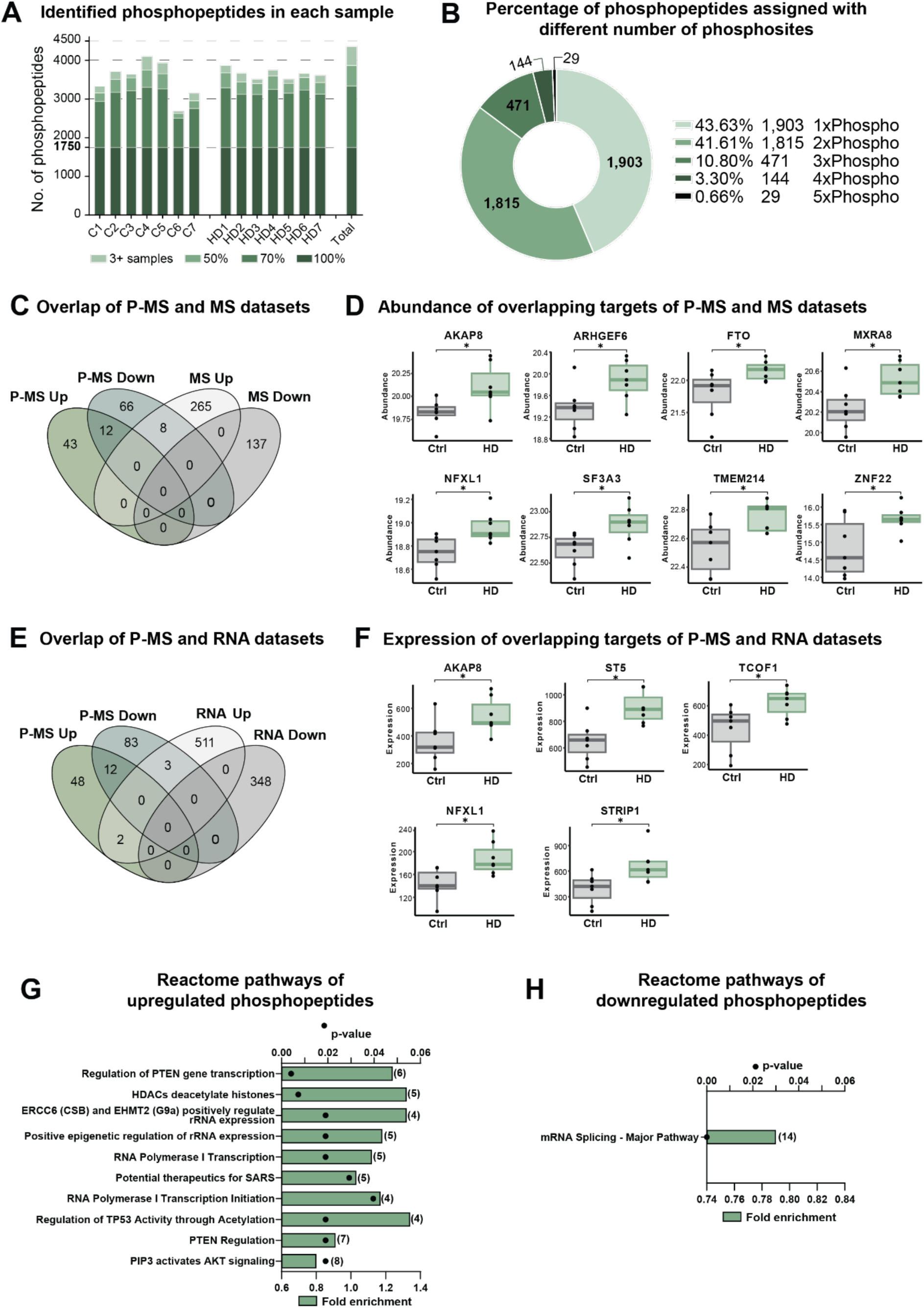
Phosphoproteome of HD-iNs. (A) Bar plots showing the number of identified phosphopeptides separately in each sample as well as in total. 1,750 phosphopeptides were present in all the samples, while 1,589, 528 and 495 phosphopeptides could be found in at least 70%, 50% and 5% of the samples. (B) Diagram presents the percentage of phosphopeptides with 1, 2, 3, 4 or 5 assigned phosphosites of all phosphopeptides. (C) Venn diagram shows the number and percentage of overlapping targets of P-MS and MS datasets in iNs. (D) Box plots of the abundance of overlapping targets of P-MS and MS datasets in n=7 Ctrl- and n=7 HD-iN samples. (E) Venn diagram shows the number and percentage of the overlapping targets of P-MS and RNA-seq datasets. (F) Box plots of RNA expression of overlapping targets of P-MS and RNA datasets in n=7 Ctrl- and n=7 HD-iN samples. (G and H) Reactome pathways using STRING-db v12.0 of upregulated phosphopeptides (G) and downregulated phosphopeptides (H). Top ten most significant terms are shown. Bar plots represent fold enrichment (strength), dots represent Benjamini-Hochberg false discovery rates (n = 7 Ctrl- and 7 HD-iN samples, FDR<0.05).

**Supplementary Figure 2.**
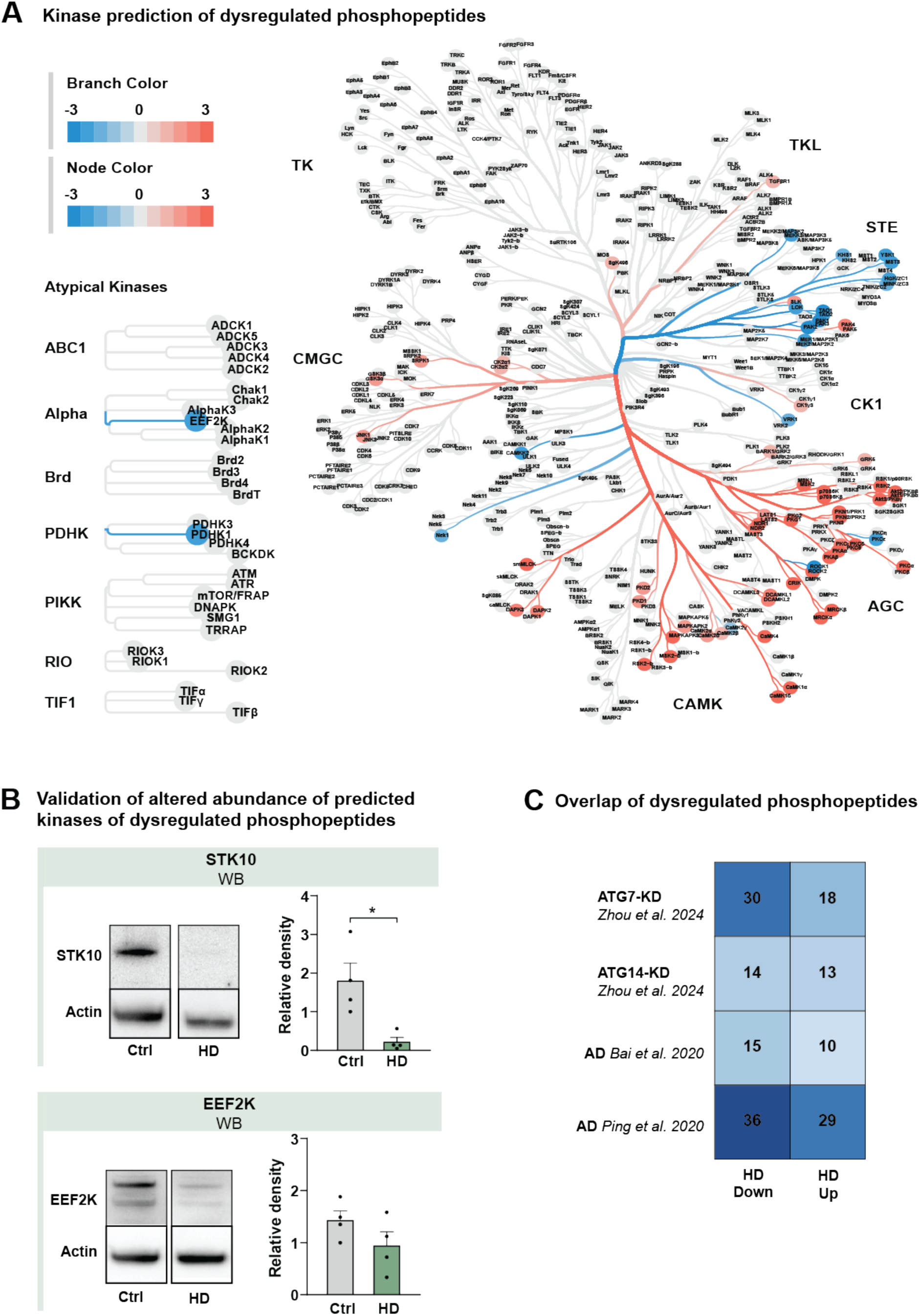
Identification of kinases of dysregulated phosphopeptides in HD-iNs and overlap of dysregulated phosphopeptides. (A) Kinase tree showing the kinases predicted to be differentially active in the HD-iNs when compared to the Ctrl-iNs. (B) Abundance of STK10 (left) and EEF2K (right) by western blot experiments in n=4 Ctrl- and n=4 HD-iN samples. Blot sections show one representative Ctrl- and HD-iN sample. Bar plots show the relative density of the proteins. (C) Heatmap of overlaps with AD and autophagy-related phosphoproteins. (B) Each dot represents one Ctrl- or one HD adult human cell line from the converted iNs, the bars show the mean value of the individual values, the error bar indicates SEM. For statistical analysis two tailed t-tests were used in all cases, * p<0.05.

**Supplementary Figure 3.**
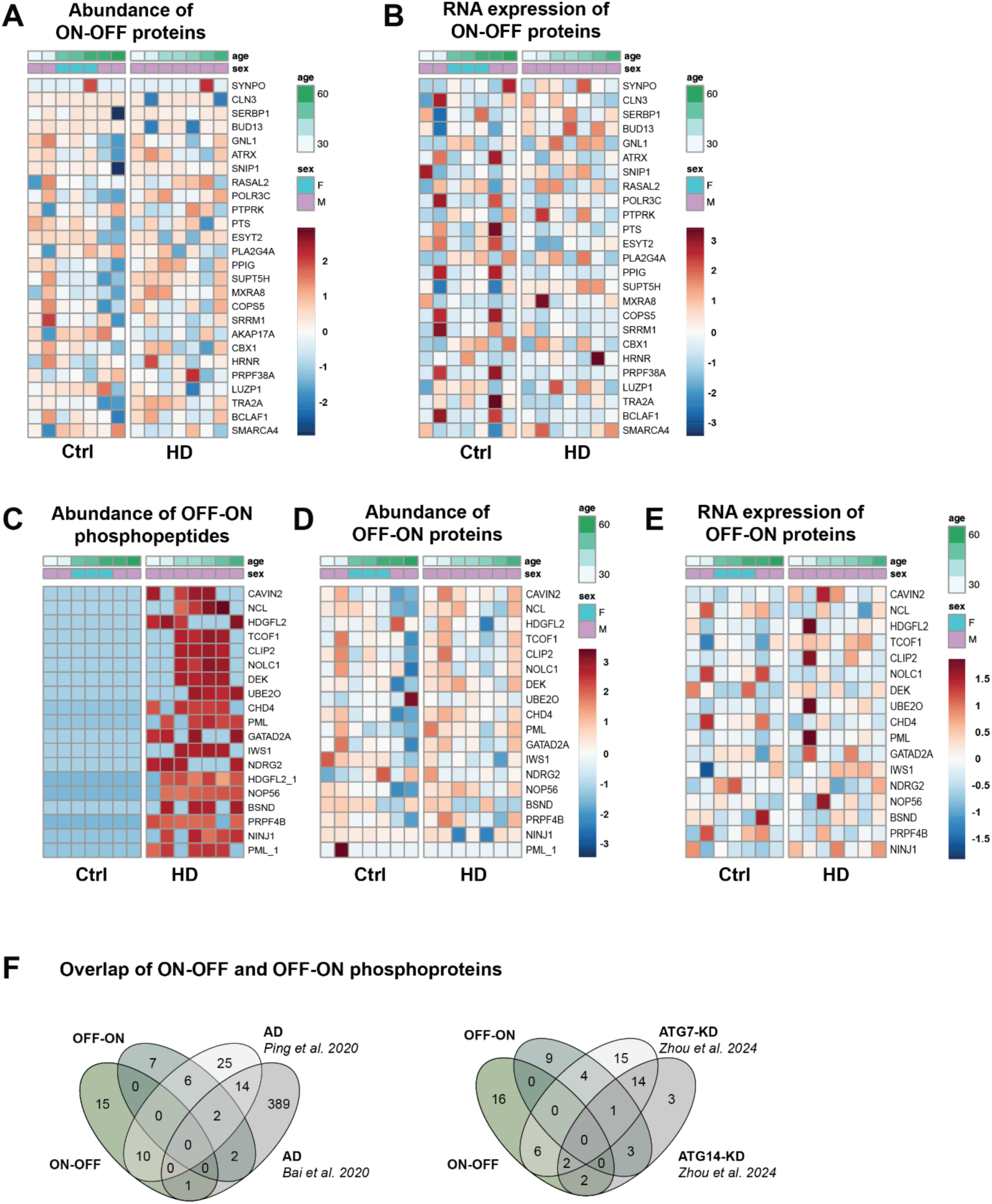
ON-OFF and OFF-ON phosphoproteins in HD-iNs. A) Heatmap of ON-OFF protein abundance in the proteomic dataset of Ctrl- and HD-iNs. (B) Heatmap of gene expression of the ON-OFF proteins in the bulk-RNAseq data of Ctrl- and HD-iNs. (C) Heatmap showing the abundance of the OFF-ON phosphopeptides in each sample. (D) Heatmap of OFF-ON protein abundance in the proteomic data of Ctrl- and HD-iNs. (E) Heatmap of gene expression of the OFF-ON proteins in the bulk-RNAseq data of Ctrl- and HD-iNs. (F) Venn diagrams showing the number of intersections of proteins associated with ON-OFF and OFF-ON phosphopeptides in the HD-iNs with those seen in (left) autophagy disruption experiments in hiPSC-iNs, and (right) studies of Alzheimer’s disease in post mortem frontal cortex samples.

**Supplementary Figure 4.**
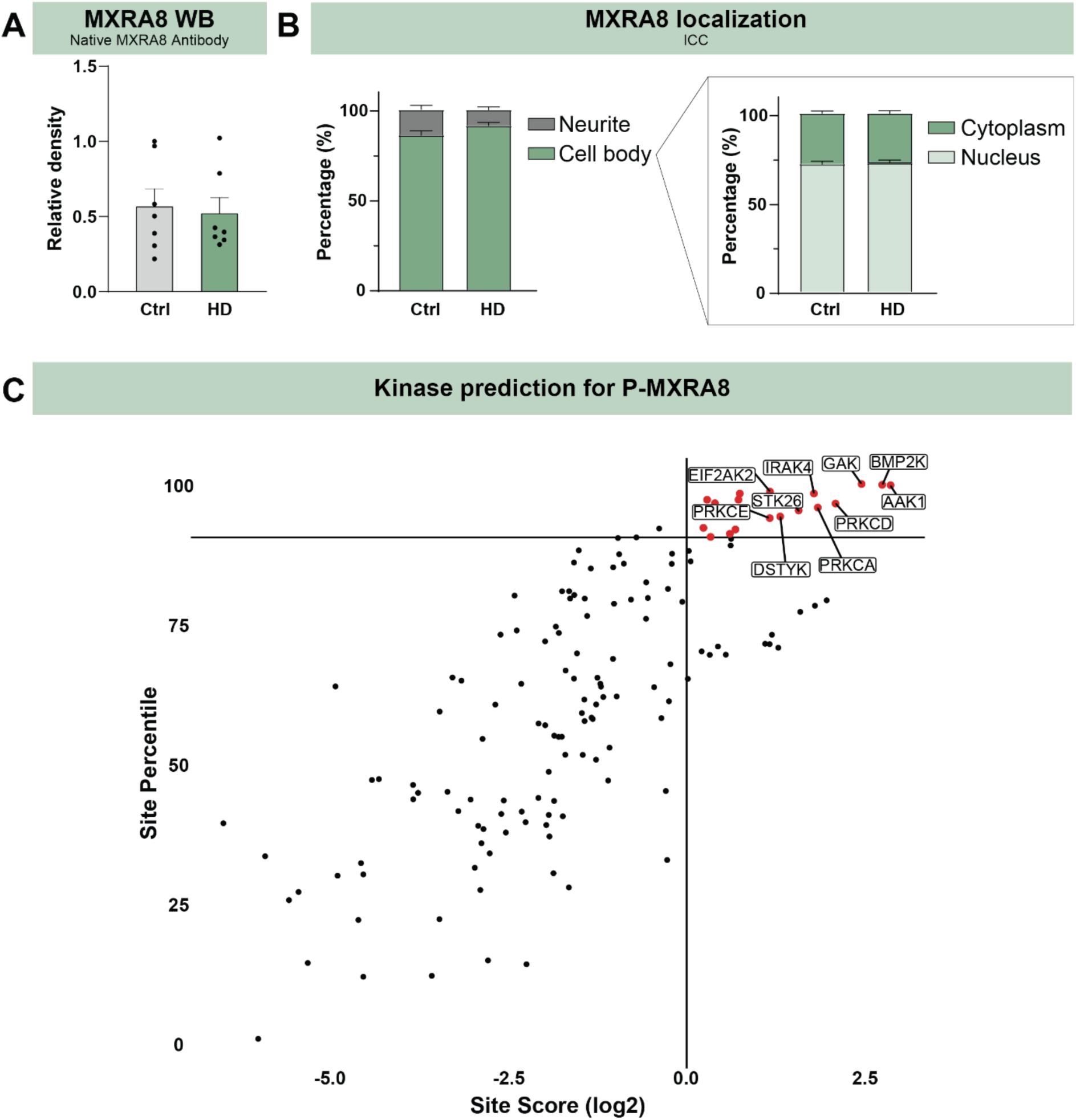
Abundance and localization of MXRA8 and its binding partners. (A) Abundance of the MXRA8 protein using native MXRA8 antibody in n=14, N=7 Ctrl- and n=14, N=7 HD-iNs. (B) Localization of MXRA8 by HCA in TAU counterstained N=7 Ctrl- and N=7 HD-iN samples. Bars on the left show the percentage of the MXRA8^+^ dots in the cell body versus in the neurites, and on the right, in the cytoplasm versus the nucleus, of Ctrl- and HD-iNs. (C) Scatter plot showing the kinase prediction results for MXRA8-pS377 from PhosphositePlus, with the Site Score on the X-axis and Site Percentile on the Y-axis. The points marked in red show the significant results, while the remaining represent the non-significant results. Any kinase with a site percentile > 90 and site score (log2) > 0, and identified in the proteomic dataset, was considered significant. (A) Each dot represents one Ctrl- or one HD-adult human cell line from the converted iNs, the bars show the mean value of the individual values, the error bar indicates SEM. For statistical analysis two tailed t-tests were used in all cases, * p<0.05.

